# New insights from 22-kHz ultrasonic vocalizations to characterize fear responses: relationship with respiration and brain oscillatory dynamics

**DOI:** 10.1101/494872

**Authors:** Maryne Dupin, Samuel Garcia, Julie Boulanger-Bertolus, Nathalie Buonviso, Anne-Marie Mouly

## Abstract

Fear behavior depends on interactions between the medial prefrontal cortex (mPFC) and the basolateral amygdala (BLA), and the expression of fear involves synchronized activity in theta and gamma oscillatory activities. In addition, freezing, the most classical measure of fear response in rodents, temporally coincides with the development of sustained 4-Hz oscillations in prefrontal– amygdala circuits. Interestingly, these oscillations were recently shown to depend on the animal’s respiratory rhythm, supporting the growing body of evidence pinpointing the influence of nasal breathing on brain rhythms. During fearful states, rats also emit 22-kHz ultrasonic vocalizations (USV) which drastically affect respiratory rhythm. However, the relationship between 22-kHz USV, respiration and brain oscillatory activities is still unknown. Yet such information is crucial for a comprehensive understanding of how the different components of fear response collectively modulate rat’s brain neural dynamics. Here we trained male rats in an odor fear conditioning task, while recording simultaneously local field potentials in BLA, mPFC and olfactory piriform cortex, together with USV calls and respiration. We show that USV calls coincide with an increase in delta and gamma power and a decrease in theta power. In addition, during USV emission in contrast to silent freezing, there is no coupling between respiratory rate and delta frequency, and the modulation of fast oscillations amplitude relative to the phase of respiration is modified. We propose that sequences of USV calls could result in a differential gating of information within the network of structures sustaining fear behavior, thus potentially modulating fear expression/memory.

**Significance Statement:** While freezing is the most frequently used measure of fear, it is only one amongst the different components of rodents’ response to threatening events. USV is another index that gives additional insight into the socioemotional status of an individual. Our study is the first to describe the effects of USV production on rat’s brain oscillatory activities in the fear neural network, and to relate some of them to changes in nasal breathing. A better knowledge of the impact of USV calls on brain neural dynamics is not only important for understanding the respective weight of the different components of fear response, but is also particularly relevant for rodent models of human neuropsychiatric disorders, for which socio-affective communication is severely impaired.

## Introduction

Fear behavior has been shown to depend on the interaction between the median prefrontal cortex (mPFC) and the basolateral amygdala (BLA), and to involve synchronized activity in theta (4–12 Hz) and gamma (30–120 Hz) frequency oscillations (Seidenbecher et al, 2003; Popa et al, 2010; Headley and Paré, 2013; Herry and Johansen, 2014; Likhtik et al, 2014; Stujenske et al, 2014; Bocchio et al, 2017). In addition, recent studies have shown that freezing, the most used index of fear response in rodents, temporally coincides with the development of sustained 4-Hz oscillations causally involved in the synchronization of spiking activity between mPFC and amygdala (Dejean et al, 2016; Karalis et al, 2016). Importantly, this slow oscillation is distinct from the theta rhythm and predicts the onset and offset of freezing. Interestingly, recent work has shown that freezing-related 4Hz oscillation in the mPFC was correlated with the animal’s respiratory rate, and that disruption of olfactory inputs to the mPFC significantly reduces the 4-Hz oscillation in the mPFC (Moberly et al, 2018). These data bring further support to the growing body of evidence showing that in addition to its impact on olfactory regions (for a review see Buonviso et al, 2006), nasal respiration also entrains oscillations in widespread brain regions including those involved in the fear network like the mPFC and amygdala (for a review see Tort et al, 2018). This suggests that the breathing rhythm, akin to slow oscillatory rhythms, could help coordinate neural activity across distant brain regions (Jensen and Colgin, 2007; Heck et al, 2017), and potentially modulate emotional/cognitive processes.

Freezing is only one amongst the different components of rodents’ response to a threatening event. In aversive situations, such as exposure to predator or footshock, rats also emit 22-kHz ultrasonic vocalizations (USV) (Schwarting and Wöhr, 2012). They indicate a negative emotional state and are associated with the termination of social behavior and the avoidance of social contacts. Their study provides a powerful tool to assess emotionality and social behavior in animal models of pathologies like autism (Wöhr and Scattoni, 2013). Surprisingly, to our knowledge no study has assessed the impact of USV production on the animal’s brain neural dynamics. Yet such information is crucial to understand how the different components of fear response collectively modulate rat’s brain neural dynamics. Indeed the emission of USV is considered as reflecting a change in emotional level and 22-kHz USV rates increase with the aversiveness of the situation, as evidenced when foot shock intensity is increased (Wöhr et al. 2005; Hegoburu et al, 2011). Importantly, 22-kHz USV emission drastically slows down the animal’s respiratory rate (Frysztak and Neafsey, 1991; Hegoburu et al, 2011; Sirotin et al, 2014; Boulanger-Bertolus et al, 2017), potentially disrupting the respiratory-related brain rhythm described above. The present study thus aimed 1) to investigate whether USV emission coincides with specific changes in oscillatory activities in the fear neural network, and 2) to assess to what extent these changes are related to changes in respiratory rate.

To do so, rats were trained in an odor fear conditioning paradigm while local field potentials (LFP) in BLA, mPFC and olfactory piriform cortex (PIR) were monitored simultaneously with USV calls, behavior and respiration, during the post-shock period. We report that USV emission temporally coincides with a significant increase in delta and gamma activities while a decrease in theta activity is observed. Some of these changes are related to respiration. Indeed, the coupling observed between respiratory rate and delta oscillation during silent freezing is lost during USV emission, and in the meantime the modulation of fast oscillations amplitude relative to the phase of respiratory cycle is modified. The present data suggest that USV calls could result in a differential gating of information within the fear neural network thus potentially modulating fear memory/expression as suggested by our observation that the amount of USV during conditioning is a good predictor of the learned fear response at retention.

## Material and Methods

### Animals

Data were obtained from twenty-three male Long Evans rats (250-270g at their arrival, Janvier Labs, France). They were housed individually at 23°C and maintained under a 12 h light–dark cycle (lights on from 8:00 am to 8:00 pm). Food and water were available ad libitum. All experiments and surgical procedures were conducted in strict accordance with the European Community Council Directive of 22nd September 2010 (2010/63/UE) and the ethics committee of University Claude Bernard Lyon 1 (CEEA-55; Approval reference DR2013-40-v2). Care was taken at all stages to minimize stress and discomfort to the animals.

### Surgery

Animals were anesthetized with Equithesin, a mixture of chloral hydrate (127 mg/kg, i.p.) and sodium pentobarbital (30 mg/kg, i.p.), and placed in a stereotaxic frame (Narishige, Japan) in a flat skull position. The level of anesthesia was held constant with regular injections of Equithesin throughout the experiment. Monopolar stainless steel recording electrodes (100 µm in diameter) were then implanted unilaterally in the left hemisphere in three brain areas: the PIR (AP: −1.8, L: +5.5mm, DV: −8 mm), the mPFC (AP: +3.0mm; L: +0.8mm; DV: −3.5mm) and the BLA (AP : −2.8mm, L : +4.9mm, DV : −7.5mm). Accurate positioning of recording electrode depth was achieved using electrophysiological characteristics of multiunit activity (BLA, mPFC) or using field potential profiles evoked by stimulation of a bipolar electrode lowered in the ipsilateral olfactory bulb (for the PIR). A reference electrode was screwed in the skull above the right parietal lobe. The three recording electrodes were connected to a telemetry transmitter (rodentPACK system, EMKA Technologies, Paris, France) fixed to the rat’s skull surface by dental acrylic cement and anchored with a surgical screw placed in the frontal bone. The animals were allowed to recover for two weeks following surgery.

### Experimental apparatus

The apparatus has been described in detail in a previous study (Hegoburu et al., 2011). It consisted of a whole body customized plethysmograph (diameter 20 cm, height 30 cm, Emka Technologies, France) placed in a sound-attenuating cage (L 60 cm, W 60 cm, H 70 cm, 56dB background noise). The plethysmograph was used to measure respiratory parameters in behaving animals. The ceiling of the plethysmograph was equipped with a tower allowing the introduction of a condenser ultrasound microphone (Avisoft-Bioacoustics CM16/CMPA, Berlin, Germany) to monitor USV emitted by the rats. The bottom of the animal chamber was equipped with a shock floor connected to a programmable Coulbourn shocker (Bilaney Consultants GmbH, Düsseldorf, Germany). Three Tygon tubing connected to a programmable custom olfactometer were inserted in the tower on the top of the plethysmograph to deliver air and odorants. Deodorized air flowed constantly through the cage (2 L/min). When programmed, an odor (McCormick Pure Peppermint; 2 L/min; 1:10 peppermint vapor to air) was introduced smoothly in the air stream through the switching of a solenoid valve (Fluid automation systems, CH-1290 Versoix, Switzerland) thus minimizing its effect on change in pressure. The bottom of the animal chamber had a port connected to a ventilation pump which could draw air out of the plethysmograph (at a rate of up to 2 L/min) thus maintaining a constant airflow that did not interact with the animal’s breathing pattern. Animal’s behavior was monitored with two video cameras on the walls of the sound-attenuating cage.

### Fear conditioning paradigm and data acquisition

After the recovery period, the animals were handled individually and placed in the experimental apparatus for 30 min each day during 3 to 4 days before the beginning of the experiments in order to familiarize them with being manipulated and connected to the telemetry transmitter.

For the conditioning session, the telemetry transmitter was plugged on the animal’s head and the rat was allowed free exploration during the first 4 min, then an odor was introduced into the cage for 20 or 30s sec, the last second of which overlapped with the delivery of a 0.4mA foot-shock. The animal received 10 odor-shock trials, with an intertrial interval of 4 min. After the last pairing, the transmitter was unplugged and the animal returned to its home cage.

### Retention test

The conditioned fear response was assessed during a retention test carried out 48h after conditioning. For the retention test, the rat was placed in the experimental cage (equipped with new visual cues and with a plastic floor to avoid contextual fear expression) and allowed a 4-min odor-free period. The CS odor was then presented five times for 20s with a 4-min intertrial interval. The animal’s freezing behavior in response to the odor was quantified. Freezing behavior was defined as a crouching posture and an absence of any visible movement except that due to breathing (Blanchard and Blanchard 1969).

### Data acquisition and preprocessing

For USV recording, *t*he ultrasound microphone was connected to a recording interface (UltraSoundGate 116 Hb, Avisoft-Bioacoustics) with the following settings: sampling rate = 214285 Hz; format = 16 bit (Wohr et al., 2005). Recordings were transferred to Avisoft SASLab Pro (version 4.2, Avisoft Bioacoustics, Berlin, Germany) and a Fast Fourier Transform (FFT) was conducted. Spectrograms were generated with an FFT-length of 512 points and a time window overlap of 87.5% (100% Frame, FlatTop window). These parameters produced a spectrogram at a frequency resolution of 419 Hz and a time resolution of 0.29 ms. The acoustic signal detection was provided by an automatic whistle tracking algorithm with a threshold of −20 dB, a minimum duration of 0.01 s and a hold time of 0.02 s. However, the accuracy of detection was verified trial by trial by an experienced user. The main parameters used in the present study were the duration as well as the peak amplitude and peak frequency of USV calls. No band pass filter has been applied during USV recording. Although a few 50-kHz USV were observed following shock delivery, in the present study we focused on 22-kHz USV.

The respiratory signal collected from the plethysmograph was amplified and sent to an acquisition card (MC-1608FS, Measurement Computing, USA; Sampling rate = 1000 Hz) for storage and offline analysis. The detection of the respiratory cycles was achieved using an algorithm described in a previous study (Roux et al., 2006). This algorithm performs two main operations: signal smoothing for noise reduction, and detection of zero-crossing points to define accurately the inspiration and expiration phase starting points. Momentary respiratory frequency was determined as the inverse of the respiratory cycle (inspiration plus expiration) duration.

The video signal collected through the two video cameras was acquired with homemade acquisition software. Offline, the animal’s freezing behavior was automatically detected using a LabView homemade software and further verified by an experimenter.

Local field potentials (LFP) were collected by telemetry via a three-channel wireless miniature transmitter (<5.2g, RodentPack EMKA Technology). LFP signals were amplified (x1000), filtered (between 0.1 and 100 Hz), digitized (sampling frequency: 1000 Hz) and stored on a computer for offline analysis.

### Data analysis

#### Data selection and experimental categories

Since the aim of the study was to assess the relationship between USV emission, respiration and brain oscillatory activity, we focused our analysis on the 1min-period following shock delivery during which USV were numerous and loud. During this period, the animal’s behavior was of two types: Freezing or Escape attempt. No other type of behavior (like grooming, exploration, quiet immobility…) was observed. Freezing was characterized as the absence of any visible movement except respiration. Escape attempt was scored when the animal showed wall climbing, running, saccadic head movements. We first noticed that while the majority of USV were emitted during freezing, a substantial amount of USV also occurred during escape behavior. This led us to distinguish four types of experimental categories: Silent Freezing, USV Freezing, Silent Escape and USV Escape (Figure 1). For each category, only segments longer than 1s were considered for further analysis.

**Figure 1:**
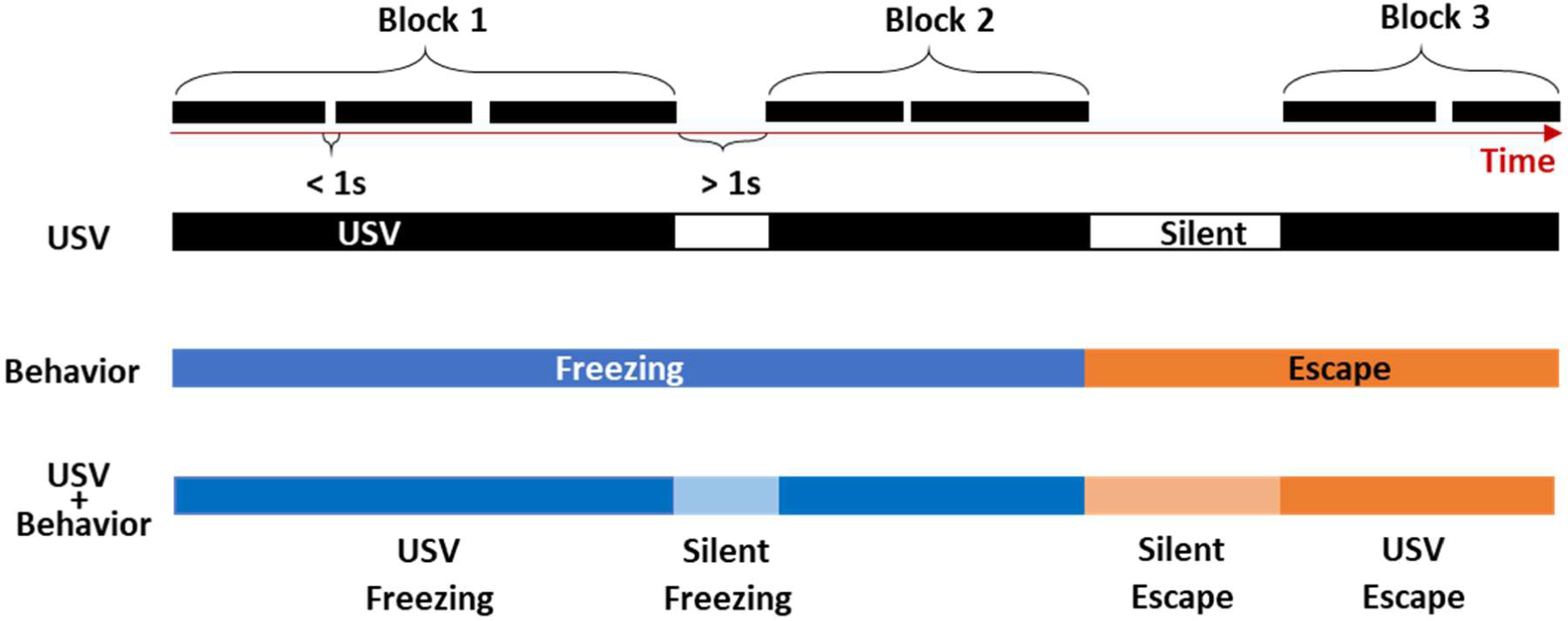
Definition of four experimental categories for data analysis. During the 1-min post-shock period, we defined blocks of USV corresponding to successive USV with less than 1s between each other. When the interval between two USV exceeded 1s, then a new block was defined (first row). The periods between USV blocks are considered as Silent periods. In parallel, the synchronized animal’s behavior (Freezing or Escape) was detected, and four different combinations were obtained: USV Freezing, USV Escape, Silent Freezing, Silent Escape. For each combination, only segments longer than 1s were considered.

#### Spectral analysis

The different data (respiration, USV, behavior, LFP signals) were synchronized offline via a TTL synchronization signal generated at the beginning of each experimental session. Once synchronized, the data were analyzed using custom-written scripts under Python.

The LFP signals were first individually inspected in order to eliminate artefacts due to signal saturation or transient signal loss. Because the average duration of individual USV calls was too short to allow proper oscillatory activity analysis (notably for the slow oscillations), we carried out the analysis on blocks of USV corresponding to successive USV with less than 1s between each other (Figure 1). As soon as the interval between two USV exceeded 1s, then a new block was defined. The periods between USV blocks are considered as Silent periods.

To extract the spectral components of the LFP signals, we used the continuous Morlet wavelet transform (Kronland-Martinet et al., 1987) instead of the classical windowed Fourier transform. Indeed the continuous wavelet transform is less susceptible to non-stationary events and offers a better time– frequency resolution. This method allowed us to segment the obtained time frequency map in periods of interest with variable durations (corresponding to the four above defined experimental categories) and average them. Four frequency bands were identified for the subsequent analyses: delta (0-5Hz), theta (5-15Hz), beta (15-40Hz) and gamma (40-80Hz). The power spectral density (PSD) of the signal was then computed, and the mean power in the different frequency bands was calculated. The values obtained for each recording site were averaged across animals.

#### Covariation of LFP slow (delta and theta) oscillatory frequency and respiratory frequency

To study frequency-frequency coupling between LFP signals and respiration, we did not use classical coherence analysis because the respiratory signal is not always sinusoidal (especially during USV calls, see Figure 5A). We therefore designed a homemade method allowing to track instantaneous frequency synchrony. To do so, for each detected respiratory cycle, the frequency was estimated as 1/cycle duration and the time course of the instantaneous respiratory frequency was extracted. In parallel, the continuous Morlet scalogram for the LFP signal was computed in our frequency band of interest (0-15Hz). At each time bin (4ms), the local maximum in the instantaneous power spectrum was extracted together with the corresponding instantaneous frequency of the LFP signal. The time course of the predominant instantaneous frequency curve of the LFP in the 0-15Hz band was then extracted. From the two times series obtained (instantaneous respiration frequency and predominant instantaneous LFP frequency), a 2D matrix histogram was built, with the respiratory frequency represented on the X-axis and the LFP frequency on Y-axis. This 2D histogram was normalized so that the total sum is 1, and point density was represented on a color scale ranging from blue to yellow as the point density increases. The existence of a coupling between respiration frequency and LFP frequency can be assumed when a high point density (ie yellow color) is observed along the diagonal of the 2D histogram (see Figure 6 for an illustration). Conversely, in the absence of coupling a non-correlated gaussian shape is observed. The two possibilities can co-occur on the same 2D histogram.

#### Modulation of LFP beta and gamma power by respiratory cycle phase

To investigate whether LFP beta and gamma amplitudes were modulated by respiration phase, we computed a so called “cycle-frequency scalogram” of the LFP signal (adapted from Roux et al, 2007). In analogy to a classical time-frequency map that computes the energy of LFP signal over time and frequency, we computed the energy of LFP signal over respiratory cycle duration and frequency. Briefly this method consists in three steps 1) Compute the continuous Morlet scalogram (time-frequency map) 2) Use detected respiratory cycle to segment this scalogram in 2 phases (inhalation, exhalation) 3) Stretch by linear interpolation each segment so that all the segments fit the same normalized template (range from 0 to 1). The result of this analysis is very similar to the classical time frequency scalogram except that it presents small time distortions locally that do not affect the instantaneous power. Since all the cycles were normalized to the same size, they were averaged together. The typical template of an individual respiratory cycle is defined as follows: inhalation from 0 to 0.4 and exhalation from 0.4 to 1. This ratio corresponds to the average value calculated over all cycles and animals. On the scalogram, LFP signal power for the different frequency bands is represented using a color scale ranging from blue to yellow as the power increases (see Figure 7B for an illustration). The maximum power in the beta and gamma bands was also extracted and represented on a curve throughout the respiratory cycle.

### Statistical analysis

All analyses were performed with Systat 13.0^®^ software. For each test, the significance level was set at p<0.05.

USV parameters (Figure 2; Duration, peak amplitude, peak frequency) were compared between USV Freezing and USV Escape using paired t-tests.

**Figure 2:**
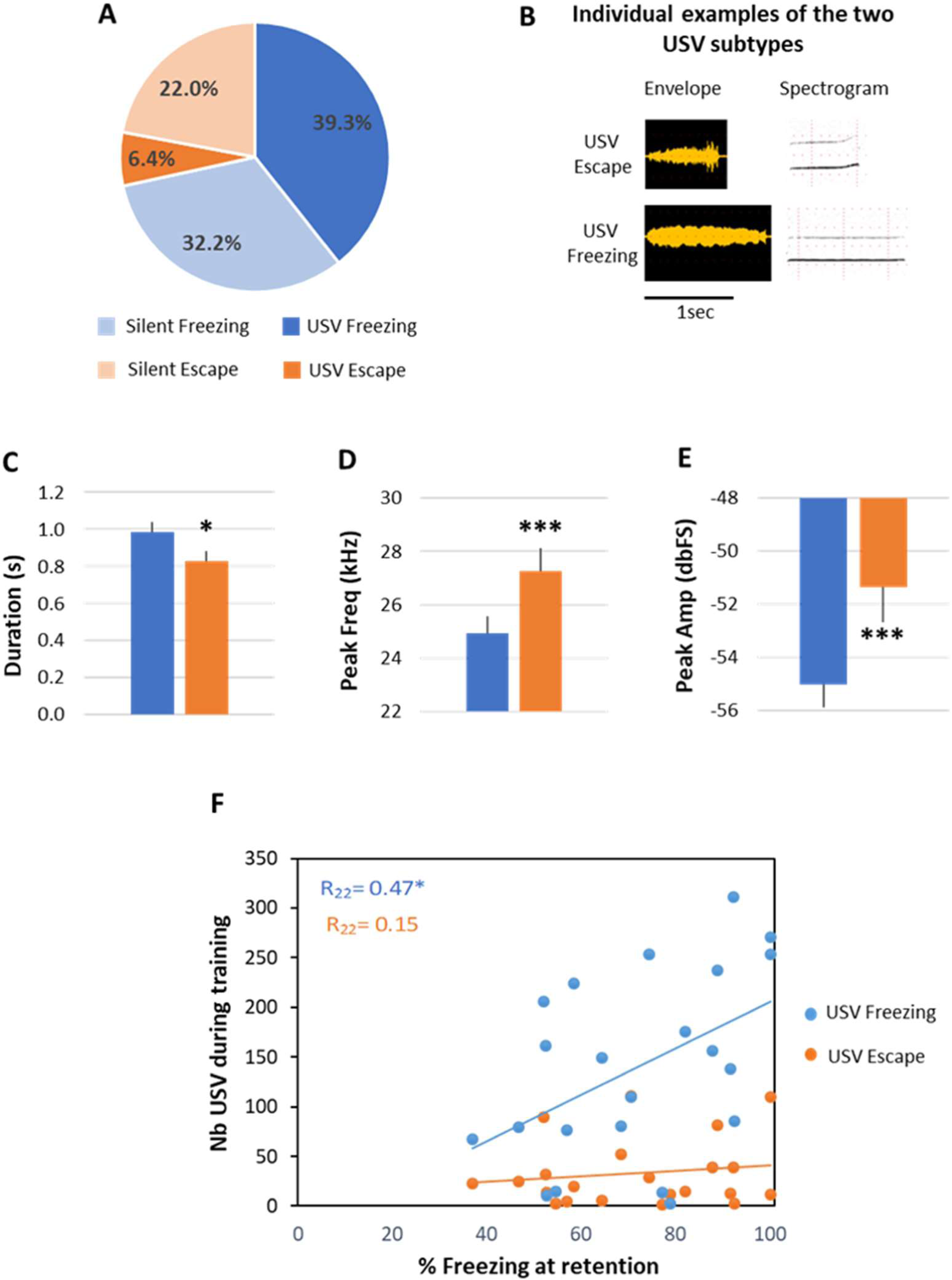
Repartition of the four defined categories and characterization of two 22-kHz USV types. A) Proportion of each category over the 1-min post-shock analysis period. B) Individual examples of raw signals (left part) and spectrograms (right part, frequency in kHz as a function of time) of the two 22-kHz USV subtypes. C) Average duration (±SEM) of the two USV subtypes. D) Average frequency (±SEM) of the two USV subtypes. E) Average intensity (±SEM) of the two USV subtypes. n=23 rats, * p<0.05, ***p<0.001. F) Correlation between the number of USV calls recorded during the 1-min post-shock period at training and the freezing score obtained during the retention test in response to the learned odor. * p<0.02

Average LFP oscillatory activity parameters (Figure 3, 4; power, peak amplitude, frequency) are calculated in the different frequency bands: delta (0–5 Hz), theta (5-15 Hz), beta (15-40 Hz) and gamma (40-80 Hz), and expressed as means ± SEM across animals. To test for differences between the four experimental categories defined above, we used two-way ANOVA for repeated measures (Factors USV and behavior) followed by paired t-tests.

**Figure 3:**
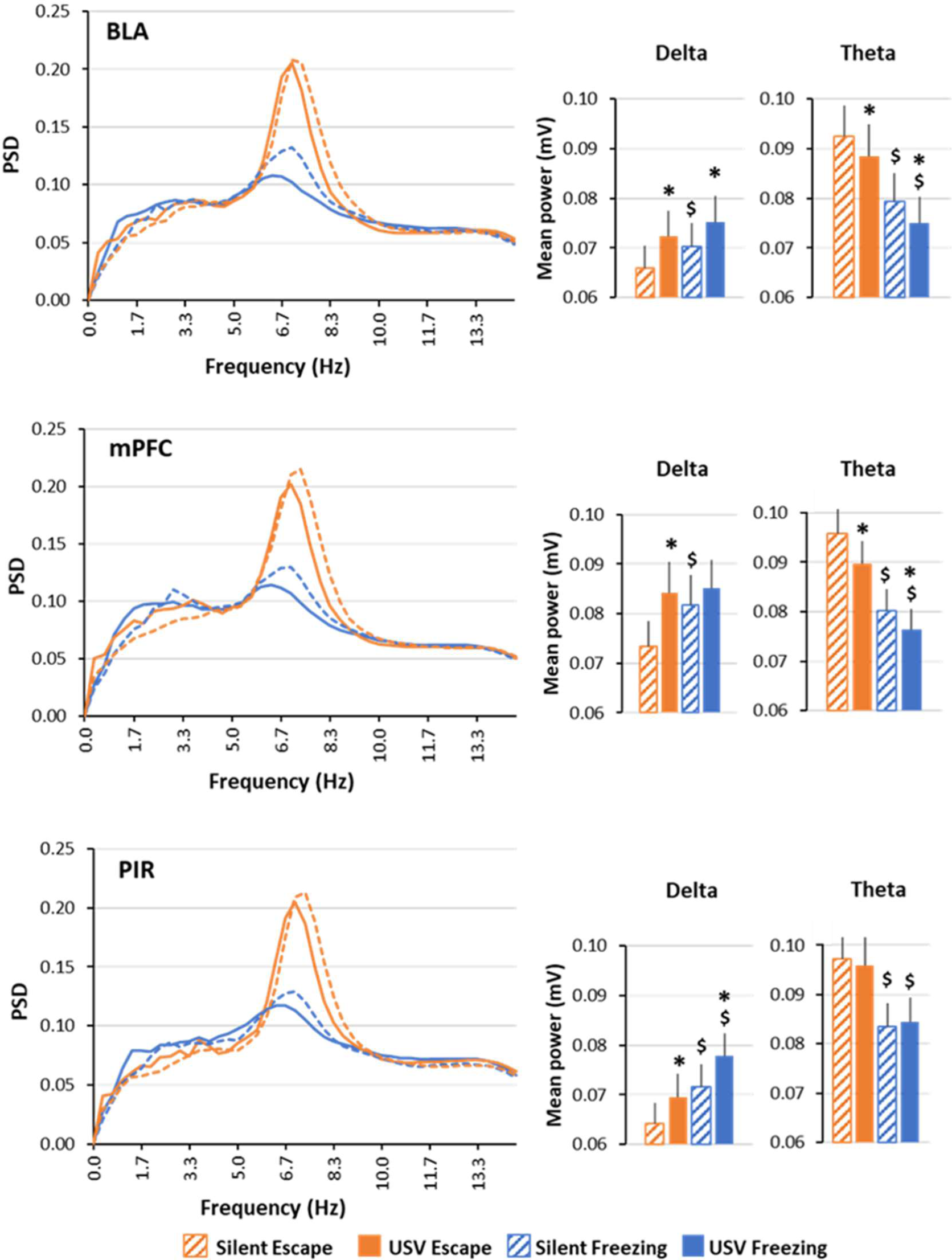
Power Spectral Density (PSD) of local field potential signals and mean power in delta (0-5Hz) and theta (5-15Hz) bands. The average PSD is represented on the left part of the figure, and the delta and theta mean power (±SEM) is represented on the right part. BLA: basolateral amygdala (n=15), mPFC: medial prefrontal cortex (n=21) and PIR: olfactory prefrontal cortex (n=20). * p<0.05, significant difference between same color-different pattern bars. $ p<0.05, significant difference between same pattern-different color bars.

**Figure 4:**
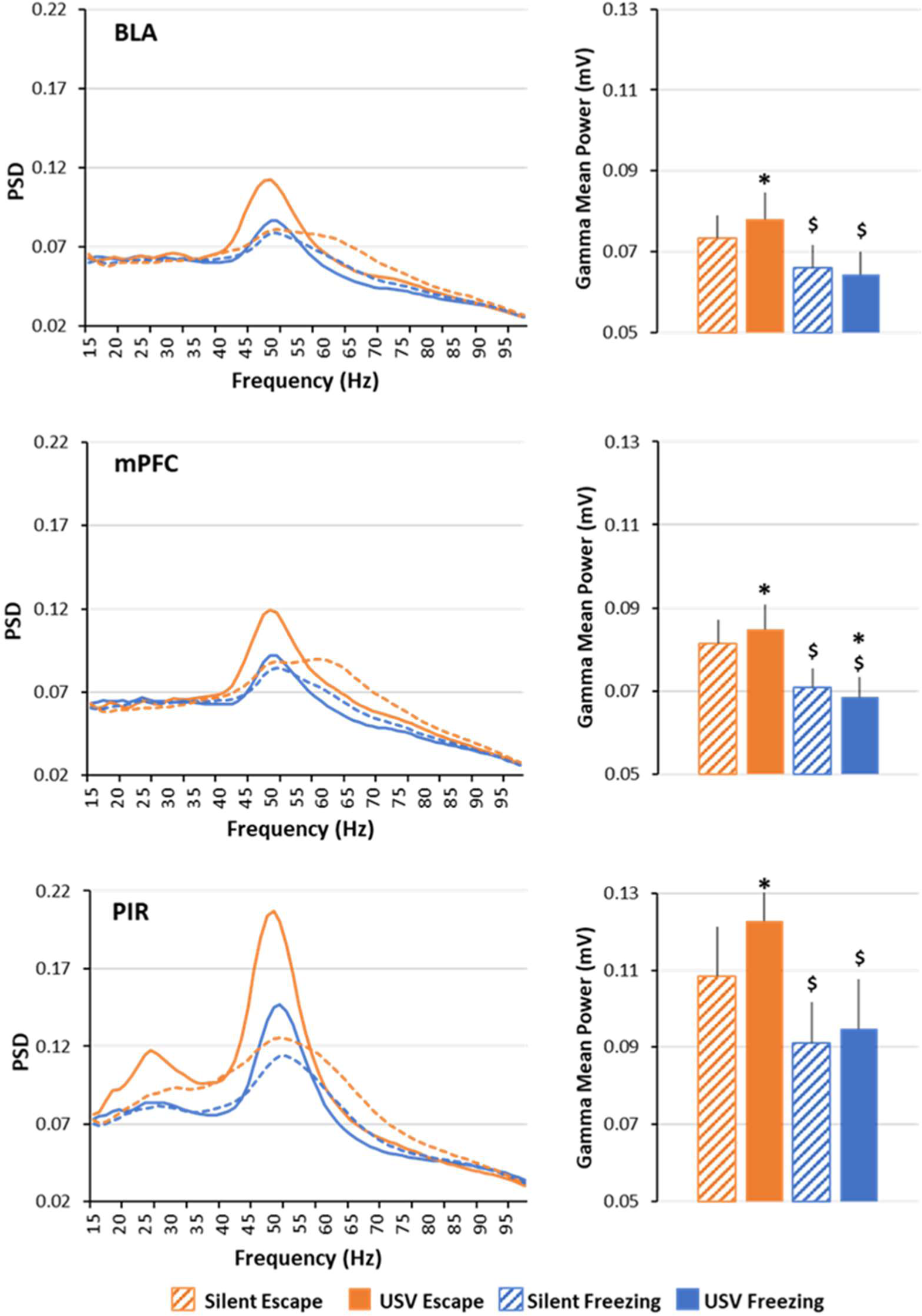
Power Spectral Density (PSD) of local field potential signals and mean power in gamma (40-80Hz) band. The average PSD is represented on the left part of the figure, and the gamma mean power (±SEM) is represented on the right part. BLA: basolateral amygdala (n=15), mPFC: medial prefrontal cortex (n=21) and PIR; olfactory piriform cortex (n=20). * p<0.05, significant difference between same color-different pattern bars. $ p<0.05, significant difference between same pattern-different color bars.

The time course of beta and gamma activity power throughout the respiratory cycle (Figure 8) was compared between silent state and USV state using a two-way ANOVA for repeated measures (Factors USV and respiratory phase).

### Histology

At the end of the experiment, the animals were sacrificed with a lethal dose of pentobarbital, their brains were removed, postfixed, cryoprotected in sucrose (20%). The brains were then sectioned (40µm coronal slices) for verification of electrodes tips by light microscopy.

## Results

### 22-kHz USV are observed during both passive and active defense responses

The 1min-period following shock delivery was analyzed for behavior, USV emission and brain oscillatory activity. During this period, the animal’s behavior was of two types: Freezing or Escape. While the majority of USV were emitted during freezing, a non-negligible amount of USV also occurred during escape attempts. This led to the distinction of four experimental categories: Silent Freezing, USV Freezing, Silent Escape and USV Escape. Figure 2A illustrates the repartition of the different categories throughout the 1min post-shock period. The animals spent 71.6% of the time in Freezing vs 28.4% in Escape. While the animals spent equivalent amounts of time in silent freezing (32.3%) compared to USV Freezing (39.3%), they spent more time in silent Escape (22%) than in USV Escape (6.4%).

We then compared the characteristics of the USV emitted during Freezing versus Escape (see Figure 2B for individual examples of the two subtypes of USV). Paired t-test comparisons revealed that call duration (Figure 2C) was significantly lower for USV emitted during escape than during Freezing (t_21_ = 2.198, p-value = 0.039), while call peak amplitude (Figure 2D) and peak frequency (Figure 2E) was significantly higher during escape than during Freezing (peak amplitude: t_21_ = −3.957, p-value = 0.001; peak frequency: t_21_ = −3.928, p-value =0.001).

Finally we assessed whether the amount of USV emitted during conditioning could predict the animal’s performance during the retention test carried out 24h later. For this, we plotted for each animal the number of USV (USV Freezing or USV Escape) emitted during the 1-min post-shock period at training against the amount of freezing exhibited in response to the learned odor during retention (Figure 2F). We showed that the number of USV Freezing was positively correlated with the amount of freezing at retention (Pearson correlation coefficient: R_22_=0.47, p<0.02) while the number of USV Escape was not (R_22_=0.15).

In summary, 22-kHz USV are emitted during both passive (Freezing) and active (Escape) defense responses. USV emitted during escape are shorter and louder than those emitted during freezing, and exhibit a higher peak frequency. In addition, the amount of USV Freezing during training was a good predictor of the animal’s learned fear response at retention.

### USV emission is associated with changes in oscillatory activity power

The local field potential signals collected in mPFC, BLA and PIR during the 1min post-shock period were analyzed separately for each recording site in the four categories described above. The main objective was to assess whether USV emission is associated with changes in oscillatory activity power compared to the silent behavioral state, and if these changes are similar across the three recording sites. Because USV are emitted during two different defense responses, we also compared oscillatory activity power between these two behavioral states (freezing versus escape).

#### Delta (0-5Hz) and theta (5-15Hz) bands mean power spectral density

For each recording site, LFP mean power spectral density was calculated per animal and averaged across animals (Figure 3, left part). Visual inspection of the spectra suggests that in each recording site, the signal power in the delta and theta bands depends on both the animal’s behavioral state (freezing vs escape) and, for a given behavioral state, the emission of USV (USV vs silent). Averaged mean power was calculated for each frequency band, in the four categories. A two-way ANOVA for repeated factors (USV and Behavior) was used to assess significant differences.

#### ***Delta band mean power*** (Figure 3, middle column)

In the three recording sites, the ANOVA revealed a significant effect of factors USV and Behavior, and no significant USVxBehavior interaction except for the mPFC (see Table 1 for all the statistical results). Paired t-test comparisons first showed that in the three structures, delta mean power was higher during silent freezing than during silent escape. In addition, in both behavioral states (except for mPFC for which the effect of USV was only significant during escape), the emission of USV was associated with a significant enhancement in delta mean power.

**Table 1:** Delta and Theta activity mean power, ANOVA analysis (upper table) and Paired t-Tests comparisons (lower table). * : significant difference

| Oscillatory Band | Structure | Factor | F (ddl) | F value | P value |
| --- | --- | --- | --- | --- | --- |
| Delta Mean power | BLA | USV | 1,13 | 17.97 | 0.001* |
|  |  | Behavior |  | 11.26 | 0.005* |
|  |  | USVxBehavior |  | 0.623 | 0.444 |
|  | mPFC | USV | 1,20 | 17.56 | <0.001* |
|  |  | Behavior |  | 5.699 | 0.027* |
|  |  | USVxBehavior |  | 9.277 | 0.006* |
|  | PIR | USV | 1,19 | 13.25 | 0.002* |
|  |  | Behavior |  | 33.22 | <0.001* |
|  |  | USVxBehavior |  | 0.146 | 0.707 |
| Theta Mean power | BLA | USV | 1,13 | 18.82 | 0.001* |
|  |  | Behavior |  | 28.43 | <0.001* |
|  |  | USVxBehavior |  | 0.01 | 0.921 |
|  | mPFC | USV | 1,20 | 27.77 | <0.001* |
|  |  | Behavior |  | 114.73 | <0.001* |
|  |  | USVxBehavior |  | 2.129 | 0.160 |
|  | PIR | USV | 1,19 | 0.005 | 0.944 |
|  |  | Behavior |  | 61.99 | <0.001* |
|  |  | USVxBehavior |  | 0.697 | 0.414 |

| Oscillatory Band | Structure | Silent Freezing vs Silent Escape | USV Freezing vs USV Escape | Silent Freezing vs USV Freezing | Silent Escape vs USV Escape |
| --- | --- | --- | --- | --- | --- |
| Delta Mean power | BLA | p=0.029* | p=0.11 | p=0.005* | p=0.006* |
|  | mPFC | p=0.001* | p=0.66 | p=0.095 | p<0.001* |
|  | PIR | p<0.001* | p=0.001* | p=0.028* | p<0.001* |
| Theta Mean power | BLA | p<0.001* | p=0.001* | p=0.017* | p<0.001* |
|  | mPFC | p<0.001* | p<0.001* | p=0.006* | p<0.001* |
|  | PIR | p<0.001* | p=0.001* | p=0.69 | p=0.49 |

#### ***Theta band mean power*** (Figure 3, right column)

In BLA and mPFC, the ANOVA revealed a significant effect of factors USV and Behavior, and no significant USVxBehavior interaction (see Table 1). In PIR, no significant effect of factor USV was observed. Paired t-test comparisons (see Table 1) showed that in the three structures, theta mean power was lower during silent freezing than during silent escape. Moreover, in BLA and mPFC, USV emission was associated with a decrease in theta mean power.

In summary, when compared to silent escape, silent freezing is characterized by a higher power of oscillatory activity in the delta band while a lower power was observed in the theta band. Importantly, in both freezing and escape states, the emission of USV coincides globally with an increase in power in the delta band in all three regions and a decrease in the theta band in BLA and mPFC.

#### Beta (15-40Hz) and gamma (40-80Hz) bands mean power spectral density

Mean power spectral density of local field potentials is represented for each recording site on Figure 4, left part. Visual inspection of the spectra suggests that in each recording site, the signal power in the gamma band depends on both the animal’s behavioral state (freezing vs escape) and, for a given behavioral state, the emission of USV (USV vs silent). Furthermore, in the PIR, the signal power in the beta band also seems to be affected during the emission of USV during escape. Averaged mean power was calculated for each frequency band, in the four categories.

#### ***Beta band mean power*** (Table 2)

No significant changes were observed in beta activity in mPFC and BLA, therefore the data are not presented in the table. In the PIR, the two-way ANOVA revealed a significant effect of USV and Behavior, and a significant USVxBehavior interaction. Paired t-tests showed that USV emission during escape induced an increase in beta mean power.

**Table 2:**
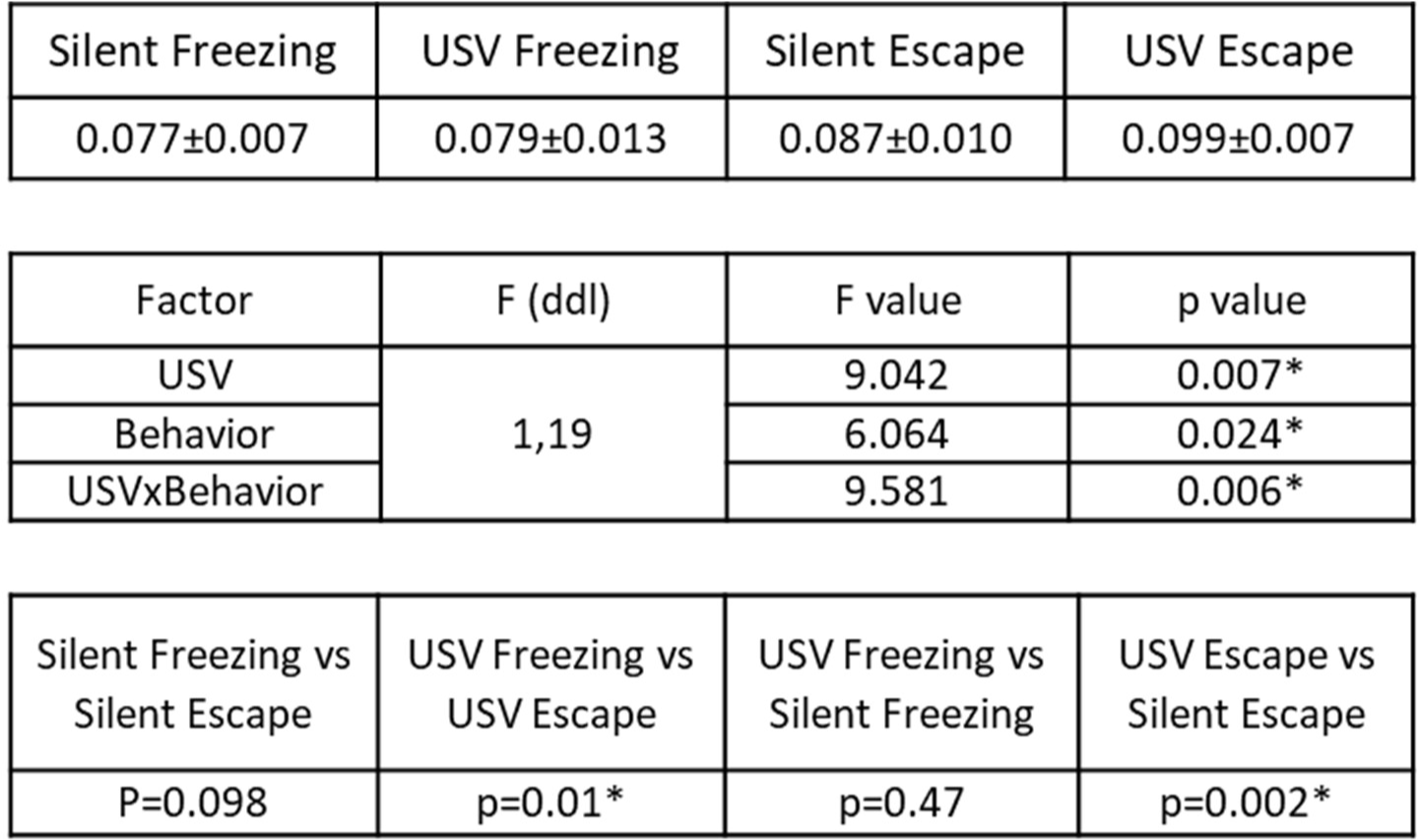
Beta band mean power (mV) +/- sem in the piriform cortex in the four experimental categories (upper table), ANOVA analysis (middle table), and Paired t-Tests comparisons (lower table). * : significant difference

#### ***Gamma band mean power*** (Figure 4, right column)

In the three recording sites, the two-way ANOVA revealed a significant effect of Behavior, and a significant USVxBehavior interaction (Table 3). Paired t-test comparisons showed that gamma mean power was lower during silent freezing than during silent escape. In addition, USV emission during escape temporally coincided with an increase in gamma mean power. Noteworthy, USV emission during both freezing and escape was associated with a narrowing of the activity towards the lower range of the band and an increase in gamma peak power (as can be seen on the power spectra of Figure 4, left column).

**Table 3:** Gamma band mean power in the three recording sites, ANOVA analysis (upper table) and Paired t-Tests comparisons (lower table). * : significant difference

| Structure | Factor | F (ddl) | F value | p value |
| --- | --- | --- | --- | --- |
| BLA | USV | 1,13 | 1.532 | 0.238 |
|  | Behavior |  | 78.54 | <0.001* |
|  | USVxBehavior |  | 6.996 | 0.020* |
| mPFC | USV | 1,20 | 0.801 | 0.381 |
|  | Behavior |  | 59.79 | <0.001* |
|  | USVxBehavior |  | 15.69 | 0.001* |
| PIR | USV | 1,19 | 6.293 | 0.021* |
|  | Behavior |  | 31.94 | <0.001* |
|  | USVxBehavior |  | 11.74 | 0.003* |

| Structure | Silent Freezing vs Silent Escape | USV Freezing vs USV Escape | USV Freezing vs Silent Freezing | USV Escape vs Silent Escape |
| --- | --- | --- | --- | --- |
| BLA | p<0.001* | p<0.001* | p=0.15 | p=0.018* |
| mPFC | p<0.001* | p<0.001* | p=0.027* | p=0.009* |
| PIR | p=0.001* | p<0.001* | p=0.413 | p=0.001* |

In summary, during silent freezing, gamma band mean power is lower than during silent escape. In addition, the emission of USV coincides with an increase in gamma band activity (mainly during escape) added to a narrowing of the activity towards the lower range of the band where the peak power is increased. Finally, in the PIR (but not in BLA or mPFC), the emission of USV during escape is associated with an increase in beta activity power.

### USV emission strongly affects instantaneous respiratory rate

As previously reported in the literature, we found that the emission of USV drastically changes the shape and frequency of the respiratory signal (see individual examples on Figure 5A). As illustrated on Figure 5B, the PDF of respiration in our four experimental categories shows that for both freezing and escape states, the emission of USV shifts the dominant respiratory frequency toward lower values, going from 6.3Hz to 1.4Hz for Escape and from 2.8Hz to 1Hz for Freezing (Figure 5B, insert). The next step of our study was then to assess to what extent the frequency of oscillatory activity in the delta and theta range followed respiratory frequency, and consequently whether USV emission has an impact on this relationship.

**Figure 5:**
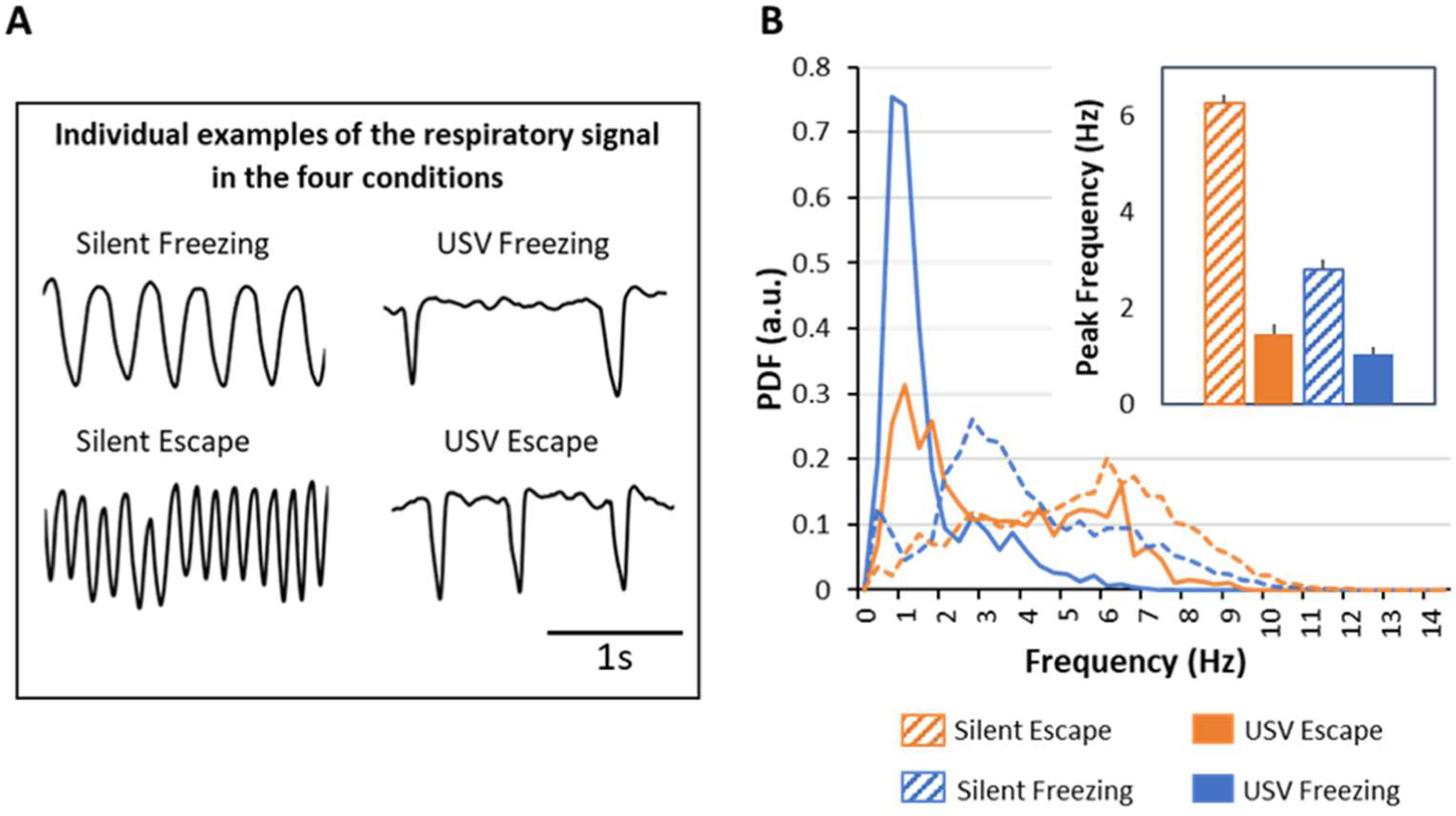
Characterization of respiratory frequency in the four experimental categories. A) Individual examples of the respiratory signal. B) Probability Distribution Function (PDF) of respiratory frequency. The distributions were obtained using a 0.33 Hz bin. Insert: Average peak frequency (± SEM) in each category (n=23 rats).

### Covariation between delta and theta oscillatory frequencies and respiratory frequency

Figure 6 (upper part) represents on the same graph the power spectral density of LFP and the PDF of respiratory signal for each experimental category. This representation highlights the fact that the respiratory frequency range delimits the range of LFP oscillatory frequency for which co-variation between the two signals frequency could be assessed.

**Figure 6:**
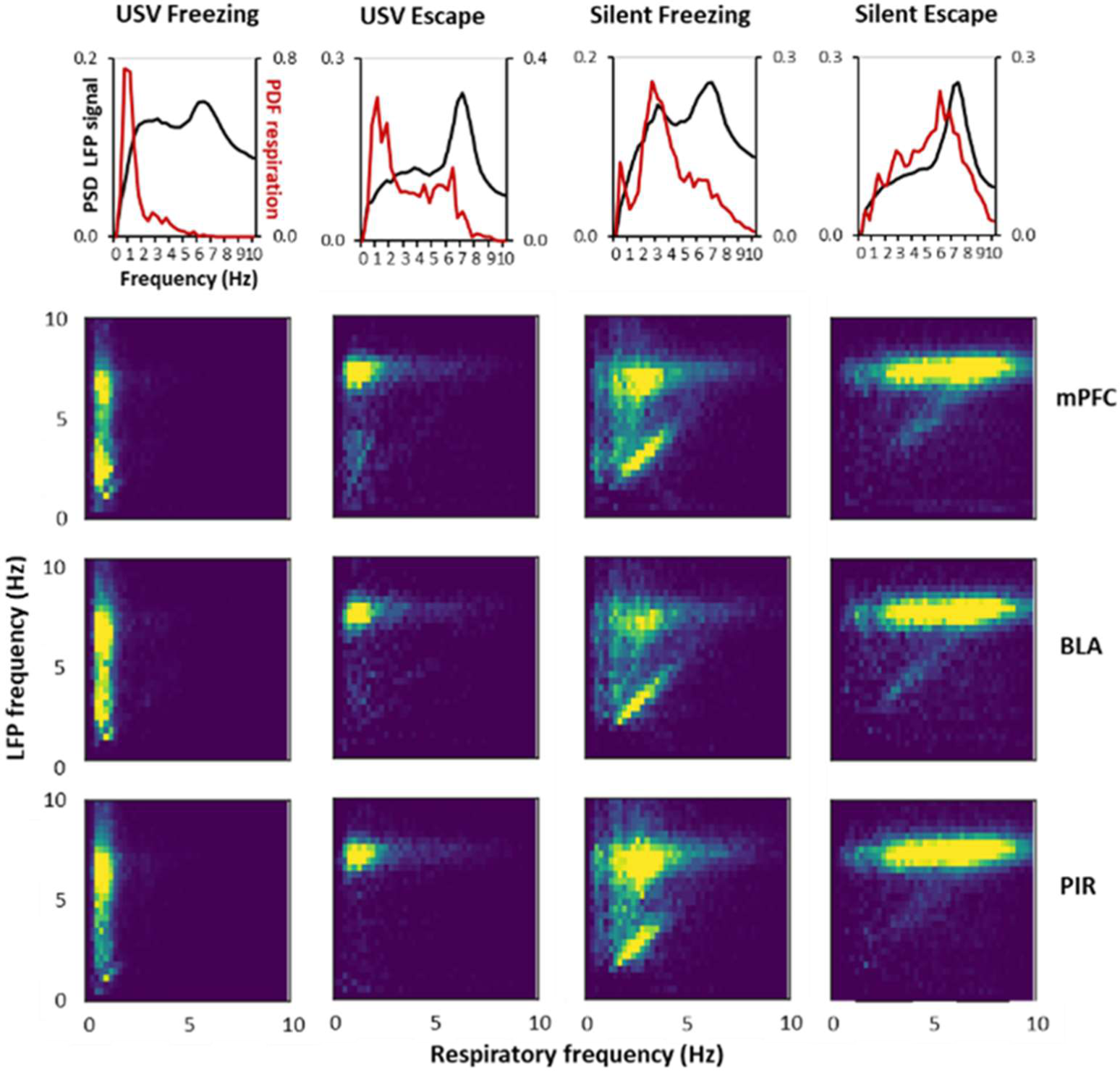
Covariation between delta and theta oscillatory frequencies and respiratory frequency. A) Each graph represents the Power Spectral Density (PSD) of local field potential (LFP) signals (Left Y axis, black curve) and the Probability Distribution Function (PDF) of respiration (Right Y axis, red curve). The graphs were obtained from LFP signals recorded in the medial prefrontal cortex in the four experimental categories (Silent Escape, USV Escape, Silent Freezing, USV Freezing). B) 2D matrix histograms obtained from LFP signals recorded in mPFC (n=21), basolateral amygdala (BLA, n=15) and olfactory piriform cortex (PIR, n=20), Y-axis represents LFP frequency and X-axis respiratory frequency. The 2D histogram is normalized so that the total sum is 1, and point density is represented on a color scale ranging from blue to yellow as the point density increases.

In each experimental category, we carried out covariation matrices (Figure 6, lower part) depicting pairwise similarities between respiratory frequency and oscillatory frequency in the delta and theta bands. During Silent Escape and USV Escape, theta activity is preferentially expressed and shows no obvious coupling with respiration: whatever the respiratory frequency, theta activity is mostly observed with a fixed frequency around 6-7Hz. During Silent Freezing, both theta and delta activities are expressed. While a clear-cut covariation is observed between delta frequency and respiratory frequency, no covariation is seen for theta frequency. During USV Freezing, both theta and delta activities are expressed with no coupling with respiratory frequency.

In summary, the only experimental category leading to a clear-cut frequency-frequency coupling between respiration and LFP signal is Silent Freezing and concerns the delta band. During USV emission, no covariation is observed between respiratory rate and delta or theta oscillatory activities.

### Modulation of beta and gamma power with the phase of the respiratory cycle

It was shown that respiration can modulate not only slow neuronal oscillations, but also beta and gamma band oscillations, which amplitude is modulated in phase with respiration (Cenier et al, 2009; Ito et al., 2014). We therefore investigated whether activity in the beta and gamma bands was modulated by the phase of the respiratory cycle in our different experimental categories, and whether USV emission has an impact on this modulation. Figure 7A illustrates an individual example of LFP signal collected in the PIR, with the corresponding respiratory signal and USV calls. A time frequency analysis carried out on the LFP signal at the level of the respiratory cycle, clearly shows that respiration modulates beta and gamma activity power, with higher beta activity power at the beginning of expiration and higher gamma activity throughout expiration. This modulation is further evidenced by the analysis illustrated on Figure 7B which represents the respiration phase-frequency map of LFP signal.

**Figure 7:**
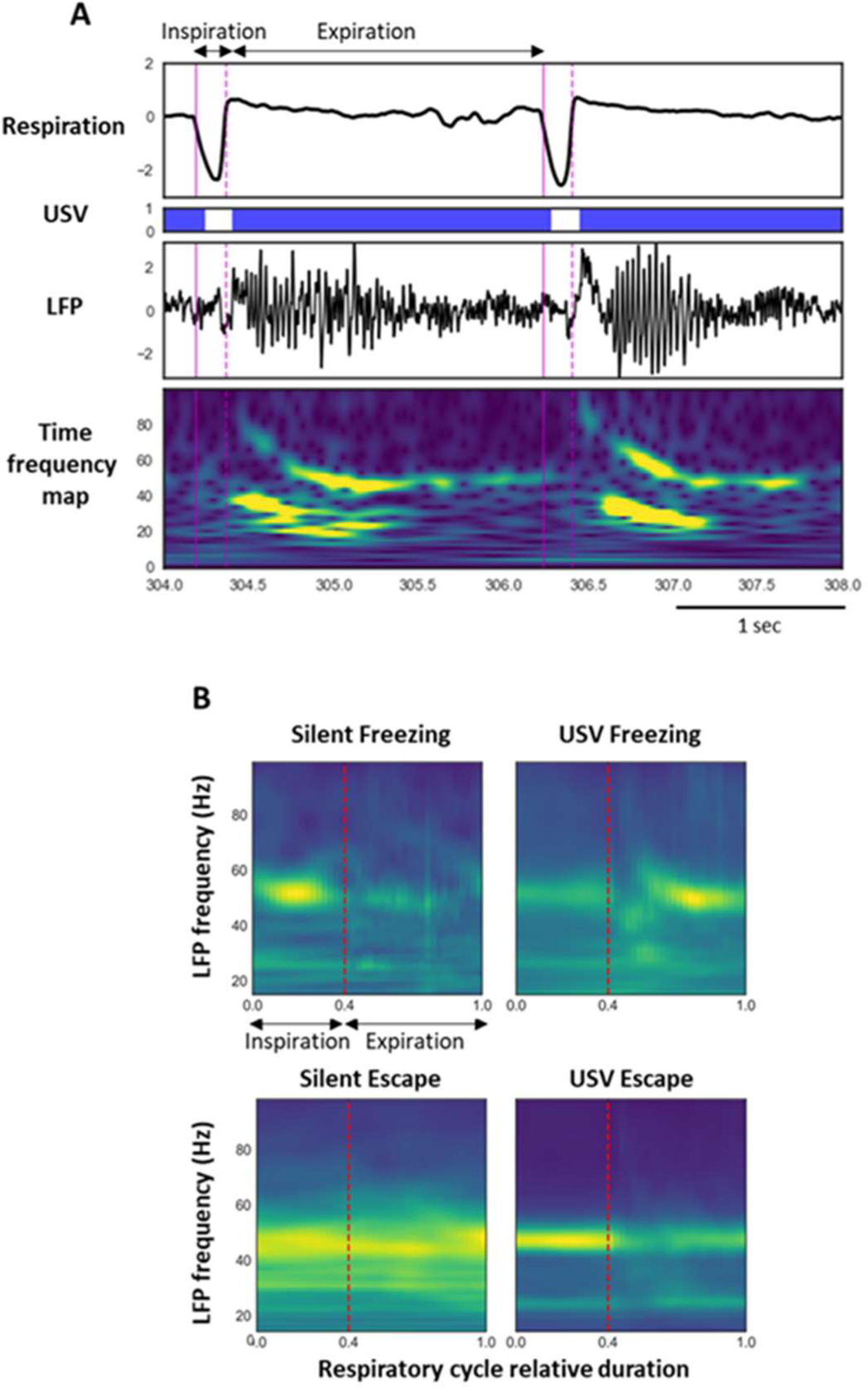
Modulation of beta and gamma power by the phase of the respiratory cycle. A) Individual traces representing from the top the respiratory signal, USV calls, raw local field potential (LFP) signal recorded in the olfactory piriform cortex and its time frequency map (Y axis: LFP signal frequency in Hz, X axis: time in ms). LFP signal power is represented using a color scale going from blue to red as the power increases. B) Average time frequency map centered on the normalized respiratory cycle. The vertical dotted line represents the transition between inspiration and expiration phase that was set at 0.4 (this value corresponds to the mean ratio between inspiration and expiration over the four experimental categories).

We carried out this analysis in the four experimental categories for each recording site. For this, in each experimental category, individual respiratory cycles were plotted on a relative duration scale (between 0 and 1), averaged per animal and further averaged across animals. The transition between inspiration and expiration was fixed at 0.4 (this value corresponds to the mean ratio between inspiration and expiration over the four experimental categories). The data illustrated on Figure 8 represent the time course of gamma and beta activity maximal power throughout the respiratory cycle. For each recording site, a two-way ANOVA for repeated factors (USV and Respiratory cycle time) was carried out separately for Freezing and Escape to compare the time course of LFP oscillatory activity power throughout the respiratory cycle, either with or without USV. In the Gamma band, during freezing (Figure 8, right part), the ANOVA revealed a significant effect of Respiratory cycle time and a significant interaction for USVxRespiratory cycle time (see Table 4). During Silent Freezing, the maximum power is observed during inspiration, while during USV Freezing, two maxima are observed, one during inspiration, and the other during the late part of expiration. Concerning escape, (Figure 8, left part), the ANOVA revealed a significant effect of Respiratory cycle time and USV (except for BLA), and a significant interaction for USVxRespiratory cycle (see Table 4). During Silent Escape, the maximum power is observed at the transition between inspiration and expiration, while during USV Escape, the maximum is shifted toward inspiration. In the Beta band in the PIR, during Freezing (Figure 8, right part), the ANOVA revealed a significant effect of Respiratory cycle time and a significant interaction for USVxRespiratory cycle time (see Table 4). During Silent Freezing, the maximum power is observed during inspiration, while during USV Freezing, the maximum is shifted toward the early part of expiration. Concerning escape (Figure 8, right part), the ANOVA revealed a significant effect of Respiratory cycle time but no effect of USV or USVxRespiratory cycle interaction (see Table 4).

**Figure 8:**
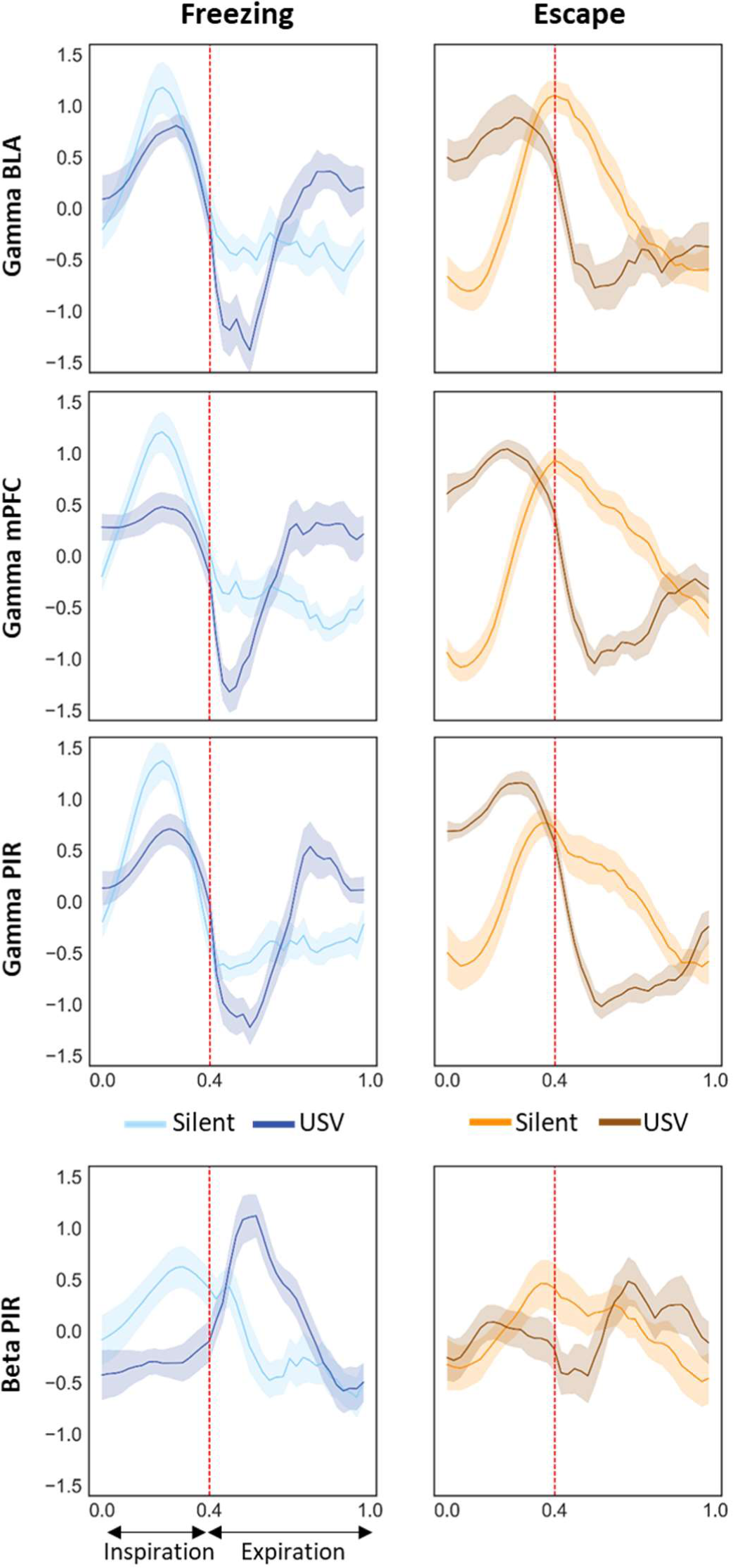
Gamma and beta activity power time course throughout the normalized respiratory cycle in the four experimental categories. Left side: Silent Freezing versus USV Freezing, right side: Silent Escape versus USV Escape. The vertical dotted line on each graph represents the transition between inspiration and expiration phase positioned at 0.4 (this value corresponds to the mean ratio between inspiration and expiration over the four experimental categories). BLA: basolateral amygdala (n=15), mPFC: medial prefrontal cortex (n=21) and PIR: olfactory piriform cortex (n=20).

**Table 4:** Gamma (upper table) and beta (lower table) bands maximum power throughout the respiratory cycle, ANOVA analysis. * : significant difference

| Gamma band | Freezing |  |  |  | Escape |  |  |
| --- | --- | --- | --- | --- | --- | --- | --- |
| Structure | Factor | F (ddl) | F value | p value | F (ddl) | F value | p value |
| BLA | Respiratory cycle time | 39,585 | 8.61 | <0.001* | 39,546 | 8.95 | <0.001* |
|  | USV | 1,15 | 0.10 | 0.761 | 1,14 | 0.10 | 0.761 |
|  | Respiratory cycle timexUSV | 39,585 | 4.28 | <0.001* | 39,546 | 4.57 | 0.051* |
| mPFC | Respiratory cycle time | 39,819 | 10.78 | <0.001* | 39,819 | 20.65 | <0.001* |
|  | USV | 1,21 | 1.05 | 0.318 | 1,21 | 6.38 | 0.020* |
|  | Respiratory cycle timexUSV | 39,819 | 4.78 | <0.001* | 39,819 | 5.62 | <0.001* |
| PIR | Respiratory cycle time | 39,780 | 14.27 | <0.001* | 39,780 | 17.62 | <0.001* |
|  | USV | 1,20 | 0.29 | 0.597 | 1,20 | 9.19 | 0.007* |
|  | Respiratory cycle timexUSV | 39,780 | 8.03 | <0.001* | 39,780 | 4.74 | <0.001* |

| Beta band | Freezing |  |  |  | Escape |  |  |
| --- | --- | --- | --- | --- | --- | --- | --- |
| Structure | Factor | F (ddl) | F value | p value | F (ddl) | F value | p value |
| PIR | Respiratory cycle time | 39,780 | 5.75 | <0.001* | 39,780 | 1.56 | 0.017* |
|  | USV | 1,20 | 0.07 | 0.80 | 1,20 | 0.76 | 0.395 |
|  | Respiratory cycle timexUSV | 39,780 | 2.73 | <0.001* | 39,780 | 1.37 | 0.066 |

In summary, gamma power in the three recording sites is strongly modulated by the phase of the respiratory cycle during both freezing and escape, although presenting slightly different patterns. The emission of USV is associated with drastic changes in the time course of this modulation. Beta power in the PIR is also modulated by the phase of the respiratory cycle during freezing and the pattern of this modulation is changed during USV emission.

## Discussion

The present study assessed for the first time the impact of 22-kHz USV production on brain dynamics in the network involved in fear expression, including the mPFC and the BLA. We report that USV emission modulates oscillatory activities differentially depending on their frequency band. Specifically, it temporally coincides with an increase in delta and gamma power, and a decrease in theta power. In addition, in the PIR, an increase in beta activity is observed. Some of these changes co-occur with USV-induced respiration changes. Indeed, during USV calls, the coupling observed between respiratory frequency and delta oscillatory frequency during silent freezing is lost, and the time course of gamma and beta power within the respiratory cycle is modified. The present data suggest that USV calls could result in a specific gating of information within the fear network, potentially modulating fear memory, as suggested by our observation that the amount of USV emitted during conditioning is a good predictor of the learned fear response at retention.

### 22-kHz USV are observed during both passive and active fear responses

22-kHz USV in rats are emitted in aversive situations such as foot-shock delivery and are considered as reflecting a negative affective state (Litvin et al, 2007; Schwarting and Wöhr, 2012; Brudzynski, 2013). We found that USV are predominantly produced during freezing, which is consistent with the literature (Brudzynski and Ocepia, 1992; Wöhr et al, 2005; Hegoburu et al, 2011; Boulanger-Bertolus et al, 2017). However, we also found that some USV are emitted during escape. Although some examples of 22-kHz USV during locomotion have been reported (Laplagne and Elias Costa, 2016; Boulanger-Bertolus et al, 2017), the characteristics of these USV remain poorly investigated. Here we show that 22-kHz USV emitted during escape are shorter and louder than those emitted during freezing and exhibit a higher peak frequency.

We also show that although these two types of USV have globally similar effects on brain oscillations, they present different relationships with performance at 48h-retention. Indeed while there is a positive correlation between the number of USV Freezing during conditioning and the amount of freezing at retention, this correlation is not found for USV Escape. This suggests that the two subtypes of 22-KHz USV reflect different aspects of fear response. For instance, USV Escape might be more related to the unconditioned response to shock, while USV Freezing are generally considered as part of the conditioned fear response (Wöhr and Schwarting, 2008).

### Freezing and Escape differentially modulate brain oscillatory activities

We first investigated whether in the absence of USV emission, the way fear is expressed is associated with different changes in brain oscillatory activities. Freezing is a passive defense response while Escape is an active response. Recent studies pinpointed that active and passive fear responses involve distinct and mutually inhibitory neurons in the central amygdala (Gozzi et al, 2010; Fadok et al, 2017). Here we show that these two response modes also differentially modulate oscillatory activity in the fear circuit. Indeed, compared to escape, freezing is characterized by a higher delta power and a lower theta and gamma power. Our data in the delta band are in line with the literature as several recent studies showed that freezing temporally coincides with the development of 4-Hz oscillations in prefrontal-amygdala circuits (Dejean et al, 2016; Karalis et al, 2016; Moberly et al, 2018). In awake animals, theta activity is known to occur preferentially during voluntary locomotor activities (Vanderwolf, 1969; Buszaki, 2002), thus explaining the increase in theta power observed here during escape. Finally, the increase in gamma power observed during escape might reflect an increased emotional level compared to freezing. Indeed, previous studies both in humans and animals have shown that gamma oscillations are enhanced during emotional situations (reviewed in Headley and Paré, 2013; Stujenske et al, 2014; Concina et al, 2018).

### 22-kHz USV emission alters respiration and is associated with changes in oscillatory activities

While several studies have investigated the neural circuit involved in USV production (reviewed in Schwarting and Wöhr, 2012), and the correlates of USV perception in the brain of conspecifics receivers (Sadananda et al, 2008; Parsana et al, 2012; Roberts and Portfors, 2015), to our knowledge no study has assessed the effect of USV production on the sender animal’s brain oscillatory activities. We showed that USV emission coincides with an increase in delta power and a decrease in theta power. In addition, an increase in gamma power is observed. The effects are globally similar for both types of USV, and in the three recording sites although small differences exist. Furthermore, an increase in beta power is specifically observed in the PIR.

Importantly, some of the changes observed during USV emission co-occurred with changes in respiratory rhythm. USV are produced during expiration upon constriction of the vocal folds resulting in an increase in subglottal pressure and a reduction of airflow through the nose (Riede, 2011; Sirotin et al, 2014). Consequently, 22-kHz USV emission induces drastic changes in the shape and frequency of the respiratory signal (Frysztak and Neafsey, 1991; Hegoburu et al, 2011; Boulanger-Bertolus et al, 2017). It is known from a long time that respiration drives oscillations time-locked to breathing cycles in the olfactory pathways (Adrian, 1942; Fontanini and Bower, 2005; Buonviso et al, 2006; Kay et al, 2009; Courtiol et al, 2011; Esclassan et al, 2012; Zelano et al, 2016) and modulates the amplitude of local beta and gamma oscillations in the olfactory bulb (Buonviso et al., 2003; Cenier et al, 2009; Rosero and Aylwin, 2011) and in the olfactory cortex (Fontanini and Bower, 2005; Zelano et al, 2016). Several recent papers have highlighted that outside its impact on olfactory regions, nasal respiration also entrains oscillations in widespread brain regions in awake rodents (Ito et al, 2014; Nguyen Chi et al, 2016; Biskamp et al, 2017; Zhong et al, 2017; Rojas-Líbano et al, 2018; Tort et al, 2018) and modulates the amplitude of fast oscillations (Ito et al, 2014; Biskamp et al, 2017; Zhong et al, 2017). Importantly a recent work has specifically investigated the link between respiration and freezing-related 4Hz oscillation in the mPFC (Moberly et al, 2018), and reported that during freezing, mice respiratory frequency is correlated with the 4-Hz oscillation in the mPFC, and that disruption of olfactory inputs to the mPFC significantly reduces the 4-Hz oscillation.

Here we show that during silent freezing, dominant frequency in the delta band covaries with respiratory frequency. In addition, we report that the activity in the gamma band for the three recording sites, and in the beta band for PIR, are modulated in phase with respiration, with greater beta and gamma power during inspiration than expiration (see summary on Figure 9). Importantly we also show that USV emission, particularly during freezing, coincides with important changes in the relationship between respiration and oscillatory activity. First, USV emission induces a deep slow-down of respiratory frequency. In parallel, the coupling between delta band dominant frequency and respiratory frequency is lost. In addition, a reorganization of beta and gamma activity power during the respiratory cycle occurs, with increased beta power in PIR during the first half of expiration phase, and increased gamma power in the three recording sites during the second half of expiration.

**Figure 9:**
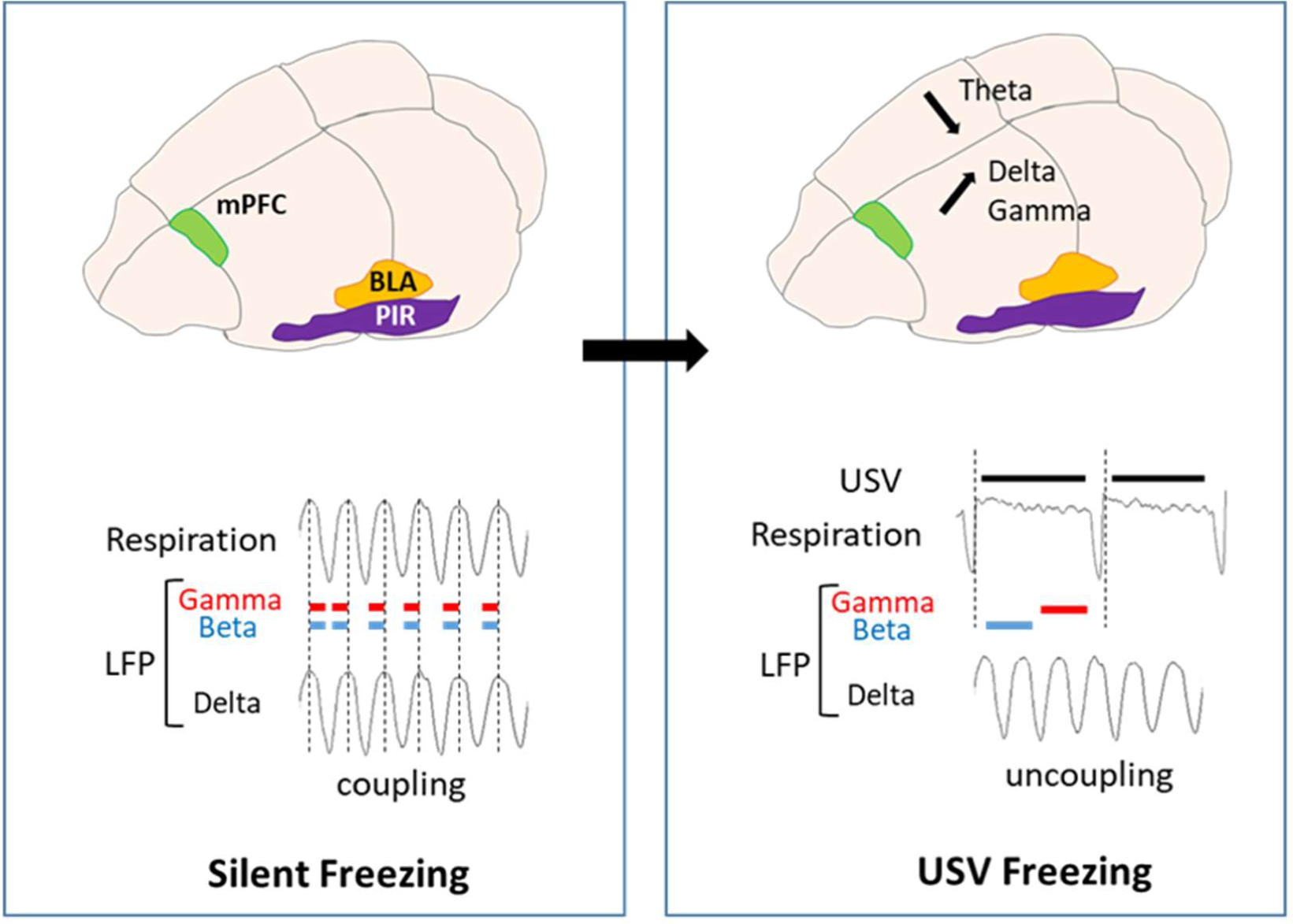
Schematic summary of the data obtained during Silent Freezing and USV Freezing. During Silent Freezing, delta frequency covaries with nasal respiratory frequency. In addition, power in the gamma band for the three recording sites, and in the beta band for the piriform cortex, is modulated in phase with respiration, with higher beta and gamma power during inspiration than expiration. USV Freezing emission coincides with a decrease in theta power and an increase in delta and gamma power. In parallel, a deep slow-down of respiratory frequency is observed, with the uncoupling between delta frequency and respiratory frequency. Furthermore, a reorganization of beta and gamma activity power during the respiratory cycle occurs, with increased beta power in the piriform cortex during the first half of expiration phase, and increased gamma power in the three recording sites during the second half of expiration. mPFC: medial prefrontal cortex, BLA: basolateral amygdala, PIR: prefrontal cortex, LFP: local field potentials

### Functional interpretation

How can we integrate the present data to the existing literature? Although additional experiments are needed to understand the functional role of USV-related changes in oscillatory activities in the fear network, tentative proposals can be made. Respiration-locked oscillations in non-olfactory regions were shown to depend on nasal airflow (Ito et al, 2014; Yanovsky et al, 2014). Indeed olfactory sensory neurons have mechanosensitive properties (Grosmaitre et al, 2007) and the signals elicited by rhythmic airflow in the nose are transmitted to the olfactory bulb and the PIR (Fontanini et al, 2003; Wu et al, 2017). The PIR has direct connections with the PFC (Clugnet and Price, 1987) and the olfactory information has unique direct access to the amygdala (McDonald, 1998). We propose that the deep slow-down of respiratory rate added to the reduction of airflow through the nose during USV calls (Riede, 2011; Sirotin et al, 2014) is responsible for the loss of coupling between nasal rhythm and delta oscillation. During USV calls, brain delta oscillations become independent of nasal respiration and their power increases. In parallel, beta and gamma activity power increases during expiration. Since we observed that the amount of USV emitted during freezing at training is correlated with the learned freezing response at retention, we suggest that the window of a USV call and its particular respiratory pattern added to its specific combination of brain oscillatory activity, might enhance plasticity at given sites of the network and ultimately strengthen long-term fear memory.

A better knowledge of the impact of USV production on brain neural dynamics is not only important for understanding the respective weight of the different components of fear response, but is also particularly relevant for rodent models of human neuropsychiatric disorders, for which socio-affective communication is severely impaired (Wöhr and Scattoni, 2013).

## Acknowledgements

This work was performed within the framework of LABEX CORTEX (ANR-11-LABX-0042) of Université de Lyon, within the program “Investissements d’Avenir” (ANR-11-IDEX-0007) operated by the French National Research Agency (ANR). The authors gratefully acknowledge Emmanuelle Courtiol and Rémi Gervais for very valuable discussions of the data and careful reading of the manuscript, and Ounsa Ben-Hellal for taking care of the animals.

Julie Boulanger-Bertolus present address: Center for Consciousness Science, Department of Anesthesiology, University of Michigan, Ann Arbor, MI, United States. The authors declare no competing financial interests.

